# Road to ruin? Urbanization increases herbivory in a tropical wildflower but does not lead to the evolution of chemical defence traits

**DOI:** 10.1101/2020.09.26.314765

**Authors:** L. Ruth Rivkin, Antonio Moura

## Abstract

Urbanization is associated with numerous changes to the biotic and abiotic environment, many of which degrade the environment and lead to a loss of biodiversity. Cities often have elevated pollution levels that harm wildlife; however, the increased concentration of some pollutants can fertilize urban plants, leading to corresponding positive effects on herbivore populations. Increases in herbivory rates may lead to natural selection for greater defence phenotypes in plants. However, evidence supporting increased herbivory leading to the evolution of plant defence in urban environments is contradictory, and entirely absent from tropical regions of the world. To address these research gaps, we evaluated herbivory on *Turnera subulata*, a common urban wildflower, along an urbanization gradient in Joao Pessoa, Brazil. We predicted that higher rates of herbivory in urban areas would lead these populations to evolve cyanogenesis, a chemical defence found in a closely related *Turnera* species. We assessed herbivory and screened for cyanogenesis in 32 populations along the urbanization gradient, quantified by the Human Footprint Index. Our results show that urbanization is significantly associated with increased herbivory rates in *T. subulata* populations. Despite elevated herbivory, we found no evidence for the evolution of cyanongenesis in any of the populations, suggesting that the fitness effects of leaf herbivory are not extreme enough to select for the evolution of plant defence in these populations. Habitat loss, predator release, and nutrient enrichment likely act together to increase the abundance of herbivorous arthropods, influencing the herbivory patterns observed in our study.

## Introduction

Urbanization is associated with numerous changes to the biotic and abiotic environment, which can have large impacts on urban biodiversity (Grimm et al., 2008; Fenoglio et al., 2020). Cities often exhibit increased pollution levels that may be harmful to urban wildlife (i.e., increased cancer risk (Sepp et al. 2019)). However, studies have indicated that increased levels of certain pollutants in the urban environment can positively benefit some species. For instance, elevated nitrogen oxides (NOx) from vehicle emissions can increase the nitrogen content and the nutritional quality of the foliage of certain species of plants (Bell et al. 2011; Honor et al, 2009; Spencer et al. 1988). This may in turn favour increased abundance of some herbivorous arthropods in urban environments (Dale & Frank, 2018; Raupp et al. 2010). Arthropod abundance in cities can also be affected through changes to their community structure (reviewed in Miles et al. 2019). Urbanization can lead to a decline in arthropod predators through habitat fragmentation and degradation, which offers a release from predation from their insect prey (Faeth et al 2005). As a result, the abundance of herbivorous arthropod species may be greater in cities relative to natural habitats.

Increased herbivorous species in cities means greater herbivory pressure on urban plants, which can compromise the growth, reproduction, and survival of plants (Mauricio & Rausher, 1997; Crawley, 1989). If this pressure sufficiently reduces plant fitness, there may have altered patterns of evolution of the defence mechanisms in urban plants (Hanley et al. 2007; Mauricio & Rausher, 1997; Miles et al., 2019; Johnson et al. 2018; Mauricio & Rausher, 1997; Züst et al. 2012). For instance, invasive wild parsnip rapidly reacquired its chemical defences when its specialist herbivore was introduced into its range (Zangerl and Berenbaum 2005; Zangerl et al. 2008). Sufficient herbivory pressure along urbanization gradients may trigger the evolution of chemical defence mechanisms in urban plant populations (Miles et al., 2019; Raupp et al. 2010), however little work has paired herbivory rates with the evolution of novel defence traits along urban gradients.

Studies that have estimated herbivory rates along urbanization gradients have shown that there is considerable spatial variation in herbivory across the urbanization gradient. However, studies differ in the results on herbivory rates between urban and rural areas; some suggest an increase in herbivory with increased urbanization (Cuevas-Reyes et al. 2013, Dale & Frank, 2017; Turrini et al. 2016) while others indicate a reduction in herbivory (Kozlov et al. 2017; Moreira et al. 2019). In addition, most studies evaluating gradients in herbivory are limited to cities in temperate regions (Miles et al. 2019). To our knowledge, the only study carried out in a tropical region found increased herbivory rates of *Solanum lycorcapurm* in habitats with greater levels urbanization (Cuevas-Reyes et al. 2013). Recent reviews on the topic (Miles et al. 2019; Rivkin et al. 2019) emphasize the urgent need for studies in tropical areas.

We assessed the effects of urbanization on herbivory and the evolution of antiherbivore defence in *Turnera subulata* in Joao Pessoa, Brazil. This species is a good system to test the effects of urbanization on herbivory and defence for several reasons. *Turnera subulata* is native to Central and South America (Schlindwein and Medeiros, 2006), and it thrives in urban habitats despite elevated pollution and habitat fragmentation. Additionally, the close relative, *T. ulmifolia*, produces the antiherbivore defence chemical hydrogen cyanide (HCN; Schappert & Shore, 1995). In cyanogenic plants, HCN is released following tissue damage, and is toxic to insect herbivores (Hughes, 1991). Cyanogenesis most likely evolved in *T. ulmifolia* if HCN as a mechanism to reduce their susceptibility to herbivores (Schappert & Shore, 1999). Although *T. subulata* has not yet been documented to express cyanogenesis (Shore & Obrist 1992), it is possible that this trait may evolve under elevated herbivory pressure. Here, we sought to answer the two questions: (1) Is there an effect of urbanization on herbivory rates on *T. subulata*? And (2) has HCN expression evolved in urban populations in response to increased herbivory pressure? Addressing these questions will help to fill the gaps in the field surrounding the effects of urbanization on species interactions in the tropics.

## Methods

### Study system

*Turnera subulata* (Passifloraceae, common names: white buttercup, sulphur alder) is a perennial herb with distylic flowers, native to Brazil and common in cities and agricultural habitats (Fig. 1). In urban areas, *T. subulata* is typically found in dense populations alongside pavement, lawns, and in dry fields. Individuals reproduce via outcrossing and is an important nectar resource for many species of bees (Schlindwein & Medeiros, 2006). *Turnera subulata* is depredated by arthropod herbivores from different feeding guilds (Cruz et al. 2019; Schappert & Shore, 1999). Most leaf herbivory is caused 23 herbivore species, although the most prevalent herbivores are lace bugs (*Gargaphia sp*., Tingidae), *Euptoieta hegesia* caterpillars, Coleoptora in the genii *Disonycha* and *Parchicola*, aphids, and leaf miners (Schappert & Shore, 1999).

**Fig. 1.**
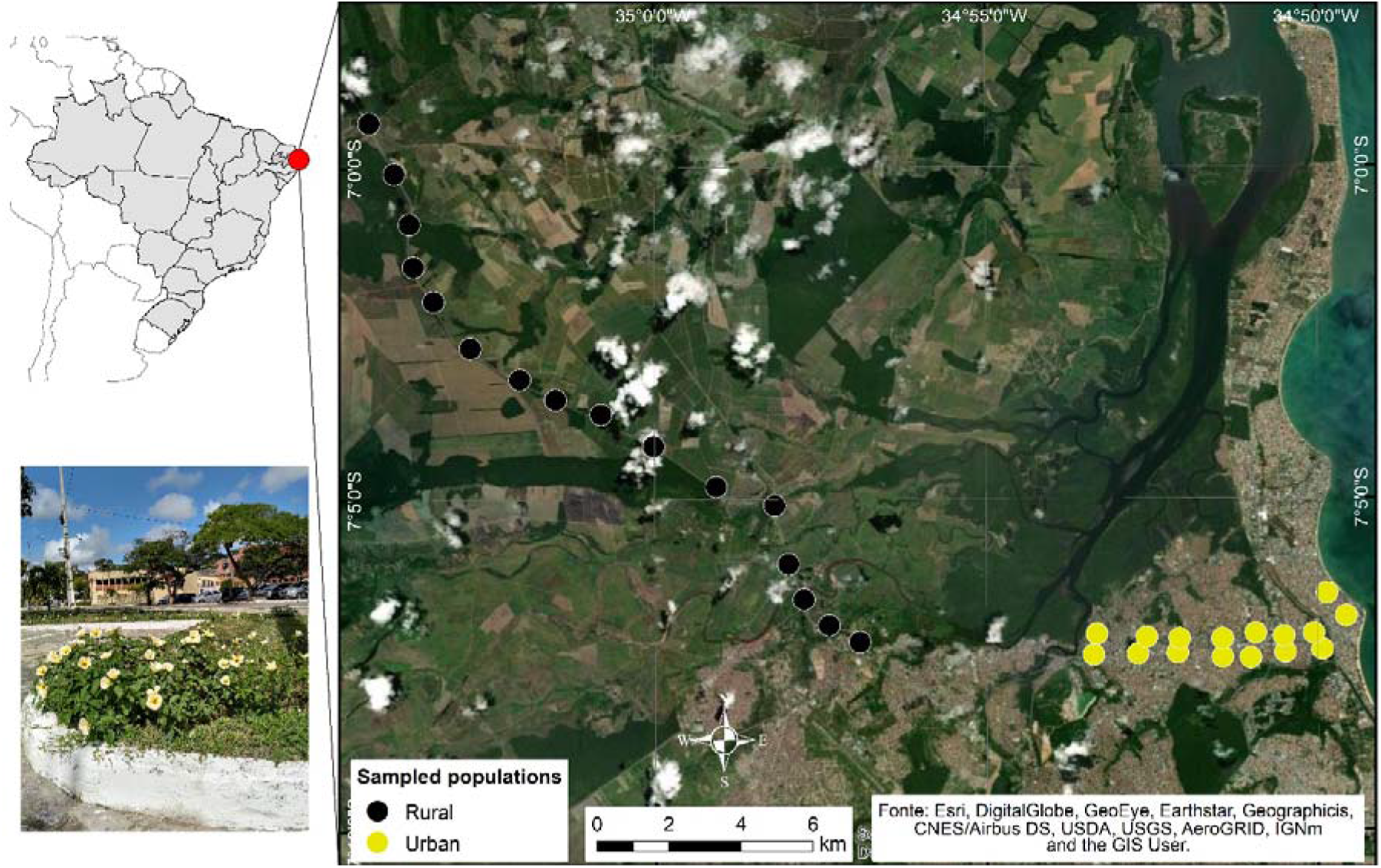
Locations of the sampled *Turnera subulata* population in Joao Pessoa in NE Brazil, demarcated by yellow (urban) and black (rural) circles. The bottom left image shows a typical urban population of *T. subulata*

### Sampling design

A total of 32 sites were selected in December 2018, along an east-west transect spanning a gradient of urbanization in the city of Joao Pessoa (>800,000 people) capital of Paraiba, north-eastern Brazil (Fig. 1). We considered a site a viable population if they consist of at least 16 *T. subulata* plants within a 100 m radius of the centre of the population. Each site in the city was at least 400 m away from the next and in the rural habitats they were at least one kilometre apart (see Fig. 1). We collected 20 plants per population, except in sites which had fewer than 20 plants (*N* = 8 sites). To avoid sampling close relatives, we sampled individual plants spaced at least 2 m apart, except in sites with fewer than 20 individuals in which case we sampled all of the plants.

We measured herbivory rate by calculating the percent area of each leaf that was damaged by herbivory. We selected the largest four leaves on up to three ramets per plant (*N*= 12 leaves per plant) and measured the amount of damage on each using the software application BioLeaf (Machado et al. 2016). This method is an effective and provide reliable measurements of leaf damage (see details in Machado et al. 2016). We calculated the per plant herbivory rate by averaging the percent leaf damage across all leaves. We also recorded whether the leaves had experienced herbivory from sap-suckling lace bugs (*Gargaphia sp*.), which discolour the leaves (Fig. S1; Guidoti et al. 2015).

We quantified the degree of urbanization at each site using the Human Footprint Index (HFI) from the GIS package generated by Venter et al. (2016a). The HFI combines human land-use variables such as increasing impervious surface and built-up areas with human demographic data to provide standardized measure of the cumulative human footprint on the terrestrial landscape (Venter et al 2016a). We used the freely available, open-source GIS software QGIS (v.3.16) to extract HFI values at 1 km2 resolution from 2009 HFI GeoTIFF dataset provided in Venter et al (2016b). We provide detailed methods on how to extract HFI values using QGIS in the supplemental materials.

### Cyanogenesis assays

We screened each plant for the presence of HCN using the Feigl-Anger assay, which use a color change reaction to indicate the presence (blue) or absence (white) of HCN (Shore & Obrist, 1992). We collected two to three young leaves from each plant, froze them at -20 oC for at least 48 hours to lyse the cell wall and induce the formation of HCN. We then followed the same protocol used by Thompson et al. (2016) to determine the frequency of cyanogenic plants in each population..

### Statistical analysis

We tested for fine-scale patterns in herbivory along the urbanization gradient by running two mixed effects regression models using R version 4.0.2 (R Core Team 2018). The first model tested for differences in the percent herbivory rate experienced by each plant averaged across all sampled leaves on a plant using a linear mixed effect model (*lmer* from the lme4 v. 1.1-23 package (Bates et al. 2015)). The second model tested for differences in the presence or absence of herbivory experienced by a plant using a generalized linear mixed effect model run on a binomial distribution (g*lmer* from lme4).

Each response variable was run against the same predictor variable: the HFI. We incorporated population ID as a random effect in each model to account for variation between plants in the population that was unrelated to urbanization. We assessed the assumptions of the models by examining the distribution of residuals, and cube-root transformed percent herbivory to better meet these assumptions. We assessed the significance of HFI on herbivory using Analysis of Variance (ANOVA) implemented with the *Anova* function in the *car* v. 3.0-6 package (Fox and Weisburg 2011) to calculate Wald chi-square test statistics with Type II sums of squares.

## Results

Herbivory rates increased with urbanization. We found that there was a significant positive effect of HFI on percent herbivory (χ^2^_1_ = 4.49, p = 0.034), where populations with the highest HFI experienced greater herbivory that populations with lower HFI (Fig. 2a). Similarly, we found there was a significant positive effect of HFI on the likelihood of herbivory (χ^2^_1_ = 10.17, p = 0.001), where urban populations were more likely to experience herbivory than nonurban populations (Fig. 2b). Two populations experience greater herbivory than others (Fig. 2), however removal of these outliers returned a similar result thus we retained them in the analysis (percent herbivory: p = 0.063, presence: p = 0.001).

**Fig. 2.**
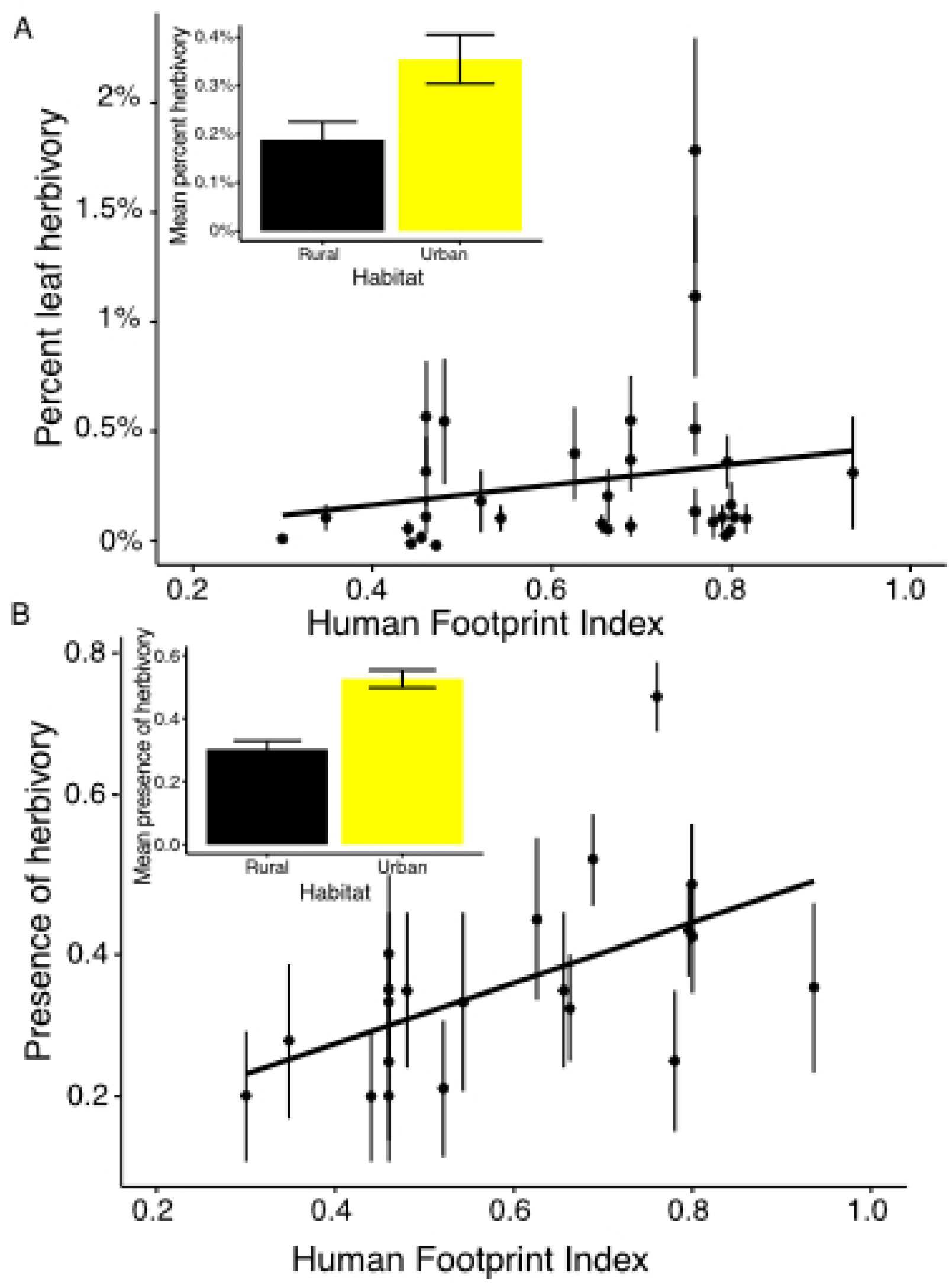
A) Percent leaf herbivory averaged across all populations increases with Human Footprint Index (p = 0.034), corresponding to greater herbivory in urban habitats (p = 0.008; inset). B) Presence of leaf herbivory averaged across all populations also increases with Human Footprint Index (p = 0.001), corresponding to increased likelihood of herbivory in urban habitats (p < 0.001; inset). Overlapping points have been jittered for clarity.

We confirmed that this pattern of herbivory was due to differences between urban and rural habitats by running the models with habitat type (urban vs rural; Fig. 1) as the predictor. Both percent herbivory (χ^2^_1_ = 7.09, p = 0.008) and presence of herbivory (χ^2^_1_ = 15.89, p < 0.001) were significantly greater in urban habitats (Fig. 2). Lastly, a two-sided t-test demonstrated that 33% of plants in urban populations exhibited evidence of lace bug herbivory whereas only 4.6% of plants in rural populations exhibited lace bug damage (t = 3.98; df = 30; p < 0.001, Figs. S2; S3).

We assayed five rural populations (97 plants) and seven urban populations (138 plants) for HCN. We detected no evidence of HCN production in any of the populations we assayed, suggesting that this chemical defence has not evolved in these populations.

## Discussion

We found that urbanization led to increased levels of herbivory on *T. subulata*, however we did not find any evidence for the evolution of HCN in any of the plants we tested. These results suggest that elevated herbivore pressure due to urbanization is insufficient to lead to the evolution of chemical defence in the populations we studied. Our results reinforce earlier findings reporting that *T. subulata* has not evolved cyanogenesis (Shore & Obrist, 1992). Arthropod herbivory appears to be the main selective driver for cyanogenesis in the closely related *T. ulmifolia* (Schappert & Shore, 1999), and this herbivore-plant interaction can drive fast evolutionary changes in plant chemical defences (Agrawal et al. 2012; Zanger et al., 2008). Nevertheless, the absence of HCN in *T. subulata* populations might be linked to more efficient mechanical protection (e.g., pilosity in leaves) or mutualistic interactions in this species that could fend off herbivory without need of HCN. Moreover, a large proportion of herbivory in urban plants were caused by lace bugs (*Gargaphia sp*.) and their feeding methods are unlikely to cause cell rupture and release of HCN, minimizing selection for this type of chemical defence.

Lace bugs feed on sap of living plants using specialized buccal apparatus that pierce the cellular tissue to extract the sap and cause serious damage in the foliage (Guidoti et al. 2015), which lead to some species being used to control population of weed plants (Baars & Heystek 2003). For instance, lace bugs are considered as one of the three most successful biological control agents against the invasive plant *Lantana camara* (Guidoti et al., 2015; Baars & Heystek 2003). Lace bugs feed gregariously on plants and cause heavy damage to the leaves (Guidoti et al. 2015) and are likely to have a strong impact on plant populations where they are abundant. In our study, we found urban plant population were more likely to exhibit lace bug damage than rural populations, suggesting that urbanization drives changes in lace bug abundance. At a community level, lace bug herbivory on *T. subulata* may be regulate urban community structure by inflicting reduction on competition within and among plant species; although speculative, this possibility deserves further study.

### City life: a road to ruin?

We found convincing evidence that urbanization is associated with elevated herbivory rates in *T. subulata* populations. This trend may be the results of increased vehicular pollution, a ubiquitous feature of urbanization. Despite its negative impacts on many organisms (Durrani et al. 2004; Rai, 2016), pollutants like NOx can increase the nitrogen content in leaves and the nutritional quality of plant foliage (Bell et al. 2011; Moreira et al. 2019; Spencer et al. 1988; Truscott et al. 2005). Increased foliage quality has been linked to high arthropod herbivore abundance in urban environment (reviewed in Miles et al 2019) because many arthropods are nitrogen limited (Dale & Frank, 2017; Jones & Leather, 2012; Raupp et al. 2010), and may be capitalizing on the increased nitrogen in roadside plants. Although we collected rural plants along a highway with high traffic load, rural plants were surrounded by fewer anthropogenic sources such as impervious surfaces, possibly reducing their exposure to pollutants.

In addition to pollution, urban habitats often exhibit reduced plant species richness due to increased habitat fragmentation (Aronson et al., 2014). Consequently, herbivory may be elevated in urban *T. subulata* sites because there are fewer species for generalist herbivores to feed upon. Lastly, rural populations may also harbour a greater diversity of herbivore predators than urban areas (Raupp et al. 2010; Turrini et al., 2016). A more diverse predatory guild in rural areas could be exerting greater top-down control compared to urban sites, reducing herbivory on plants in rural habitats. A number of studies suggest urbanization is associate with a reduction of predator and parasitoid species, leading to a release form predation in lower-guild species (Burkman & Gardner, 2014; Denys & Schmidt 1998; McIntyre 2000; Rocha & Fellowes, 2020). The decline in richness at high trophic levels in urban environment appears to be a pervasive trend and has been associated with increased herbivory in other systems (Rivkin et al 2020; Turrini et al. 2016; Raupp et al. 2010; Cuevas-Reyes et al. 2013; McIntyre 2000).

A complex relationship between abiotic environmental factors and biotic interactions, exists in urban system, and how these factors jointly influence the strength of the trophic effects on food webs is yet not well understood (El-Sabaawi 2018; Lagucki et al. 2017; Turrini et al. 2016). For example, habitat fragmentation and increased barriers to dispersal (e.g., roads) in cities reduce connectivity, and may lead to lowered abundance and diversity natural enemies to herbivores (Peralta et al. 2011; Rocha & Fellowes, 2020). Spatial distribution and patches size of plants could affect the abundance of arthropod species, leading to altered patterns of herbivory in cities (Fenoglio et al., 2020; Peralta et al. 2011; Raupp et al., 2010). We suggest that the extensive human footprint found in cities causes significant shifts in arthropod communities, releasing populations of herbivorous from top-down control, and facilitating their increased abundance in cities. However, we also found that this herbivory pressure was insufficient for *T. subulate* to evolve chemical defence to resist herbivory, thus the role of herbivory as an ultimate driver of plant defence in urban environments remains unclear.

